# Compositional analysis of microbiome data using the linear decomposition model (LDM)

**DOI:** 10.1101/2023.05.26.542540

**Authors:** Yi-Juan Hu, Glen A. Satten

## Abstract

**Summary:** There are compelling reasons to test compositional hypotheses about microbiome data. We present here LDM-clr, an extension of our linear decomposition model (LDM) approach to allow fitting linear models to centered-log-ratio-transformed taxa count data. As LDM-clr is implemented within the existing LDM program, it enjoys all the features supported by LDM, including a compositional analysis of differential abundance at both the taxon and community levels, while allowing for a wide range of covariates and study designs for either association or mediation analysis.

**Availability and Implementation:** LDM-clr has been added to the R package LDM, which is available on GitHub at https://github.com/yijuanhu/LDM.

**Supplementary information:** Supplementary data are available at Bioinformatics online.

## Introduction

Microbiome association studies detect taxa that are most strongly associated with the trait of interest (e.g., a clinical outcome or environmental factor), which are then used as microbial biomarkers for prognosis and diagnosis of a disease or microbial targets (e.g., pathogenic or probiotic bacteria) for drug discovery. Data derived from 16S amplicon or metagenomic sequencing are generally summarized in a taxa count table. However, since the total sample read count (library size) is an experimental artifact, only the relative abundances of taxa can be measured, not their absolute abundances. Consequently, microbiome data are compositional, meaning that a change in one taxon’s relative abundance necessitates a counterbalancing change in the relative abundance of (at least one) other taxa. The *compositional* null hypothesis tested for a taxon posits that the ratios of taxon relative abundances are constant across varying trait values. This hypothesis is frequently expressed in terms of parameters describing log-transformed data; the centered log ratio (clr), which centers the log relative abundance of each taxon by the sample mean log relative abundance of all taxa is most frequently used [1]. Since zero count data cannot be log-transformed, it is customary to add a pseudocount, usually 0.5 or 1, to the zeros or all entries in the taxa count table [1, 2].

We previously developed the linear decomposition model (LDM) [3–8] for testing hypotheses about the microbiome. Initially, LDM was designed for testing the null hypothesis that the relative abundance at a given taxon remains constant as the trait changes; this hypothesis could also be tested using arcsin-root-transformed relative abundance data. Later, LDM was extended to accommodate presence-absence data [5]. LDM is based on a linear model that regresses taxon data on traits/covariates, using a permutation procedure for inference. The linear model and permutation-based inference render LDM highly versatile, making it suitable for a wide variety of analyses (e.g., association, mediation), taxon data scales (e.g., relative abundance or presence-absence), trait types (e.g., continuous, discrete, multivariate, and time-to-event), and sample structures (e.g., matched sets, clustered data). Further, LDM gives a unified taxon- and community-level analysis. A natural extension of LDM for compositional analysis would be to incorporate clr-transformed taxon data. Because the parameters tested in the compositional hypothesis involve more than one taxon, some changes to LDM are required, as described herein. The resulting tests still inherit the aforementioned features of LDM, as also summarized in Figure S1. In this paper, we present LDM-clr, an extension of LDM that allows analysis of clr data.

LinDA [9], a recently published method for compositional analysis of microbiome data, also relies on a linear model for clr-transformed data. It has demonstrated better FDR control and higher sensitivity than many existing compositional methods, including ANCOM-BC [2], ALDEx2 [10], metagenomeSeq [11], and MaAsLin2 [12]. However, LinDA bases its inference on asymptotic distributions, which may not be appropriate for small sample sizes, small numbers of taxa, and complex data structures. Furthermore, LinDA is not applicable for multivariate traits or time-to-event (survival) outcomes and cannot be used for testing community-level hypotheses.

## Methods

Let *Y*_*ij*_ denote the number of sequence reads from sample *i* assigned to taxon *j* (*j* = 1, 2, …, *J*), *X*_*i*_ represent a (possibly multivariate) trait of interest we wish to test, and *Z*_*i*_ indicate a set of potentially confounding covariates we wish to control for. Optimally, we would like to test hypotheses about how the absolute abundance of each taxon depends on the covariates. However, the information on absolute abundance is typically not available; even if the total bacterial counts are measured separately (e.g., using flow cytometry), the biases inherent in microbiome data make these values unreliable [13]. If instead we formulate a log-linear model for the *observed* counts *Y*_*ij*_, we must write

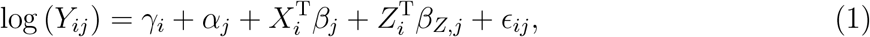

where *γ*_*i*_ is an unknown, sample-specific size factor that encodes information on the ratio of observed to true counts, *β*_*j*_ is the association parameter of interest, *α*_*j*_ and *β*_*Z,j*_ are nuisance parameters and *ε*_*ij*_ is a mean-zero error. The intercept *α*_*j*_ is the sum of a term corresponding to the log relative abundance of taxon *j* when *Z*_*i*_ and *X*_*i*_ are both zero and an unknown bias factor due to experimental bias [13–15]. The standard LDM cannot be used to fit (1) because of the *γ*_*i*_s.

The compositional approach considers the *γ*_*i*_s to be unestimable, and instead eliminates the *γ*_*i*_s by writing a linear model for some form of log-ratio data. Here we consider the clr, given by 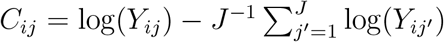 Following from (1), we have

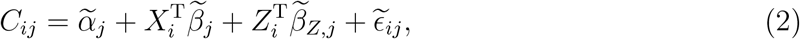

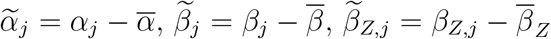 and _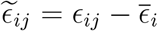_ where 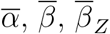, and 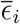 are the averages of *α*_*j*_, *β*_*j*_, *β*_*Z,j*_, and *ε*_*ij*_ over all taxa, respectively. Model (2) *can* be fit using LDM, but the parameters are not the same as those in (1) as they are centered by their means over all taxa, which makes inference on individual taxa difficult. In particular, the usual procedure in LDM of testing 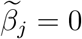 in (2) will not correspond to an easily interpretable result about the association between covariates *X*_*i*_ and taxon *j*.

Denote the least-square estimators of 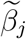and 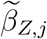 obtained using the LDM by 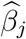 and 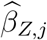 respectively. In order to obtain inference on individual taxa, we follow the approach in LOCOM [15] and assume that the majority (i.e., more than half) of the taxa are null taxa. This assumption has been frequently adopted in compositional methods (e.g., [15–17]). With this assumption, we expect that the median of the 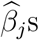 denoted by 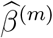 will correspond to a taxon that is null. If this is the case, then we expect 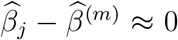 if taxon *j* is also null. When 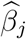 is a vector, the median is calculated in an element-wise manner.

If the number of taxa *J* is large, we can treat 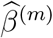 as fixed; if we subtract 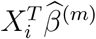 from both sides of (2) then the estimated coefficients are automatically 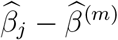 Then, LDM-clr performs permutation tests based on the proportion of permutation replicates in which the *F* statistic

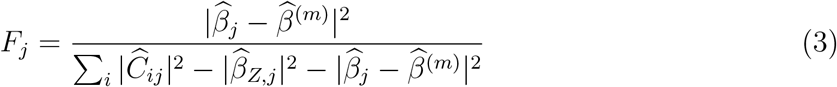

exceeds the observed value, where 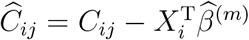

When the number of taxa *J* is small, 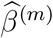 may contribute considerable variability. In this case, we still use (3), but use the value of 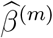 and hence 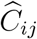 calculated for each permutation replicate. Thus, the total sum-of-squares in the denominator of (3) is different for each replicate; the “missing” variability corresponds to the variability in 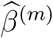 Similarly, the global test in LDM-clr sums numerators and denominators in (3) separately. The change in total variability 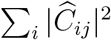 across replicates does not affect the validity of the test because it is permutation-based. The flag `adjust.bias=TRUE` tells LDM-clr to use 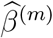 from each replicate the *F* statistic. In our simulations, this adjustment was only necessary for *J* < 100.

Like LOCOM and LDM-clr, LinDA [9] also bases inference on 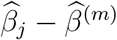 although LinDA refers to 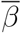 as a “bias” rather than recognizing that the regression coefficients in models (1) and (2) are simply different. LinDA estimates the “bias” 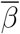 using the mode of the 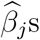 leading to the odd conclusion (in their framework) that the mode be used to estimate the mean of the *β*_*j*_ values. Still, in our framework, using the mode in place of the median when calculating 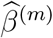 is attractive in some settings, and is offered as an option in LDM-clr.

## Results

### Simulation studies

We evaluated LDM-clr using the same set of simulation settings used to assess LinDA [9]: S0C0 (a binary trait), S0C1 (a continuous trait), S0C2 (a binary trait with two confounders), S1C0 (zero-inflated absolute abundances), S2C0 (correlated absolute abundances.), S3C0 (Gamma abundance distribution), S4C0 (small number of taxa), S5C0 (small sample size), S6C0 (10-fold difference in library size), S7C0 (Negative-Binomial abundance distribution), S8.1C0 (pre-treatment and post-treatment comparison), and S8.2C0 (replicate sampling). We applied LDM-clr using both the mode and median as 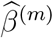 referred to as LDM-mode and LDM-median, respectively, for testing individual taxa and the entire community. Additionally, we applied LinDA for testing individual taxa only.

First, we successfully replicated the results of LinDA as reported in [9]. In S4C0 (Figure 1, left), where LinDA yielded inflated FDR due to a small number of taxa, LDM-mode and LDM-median controlled the FDR as they adjusted for the uncertainty in the value of 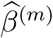 In S8.2 (Figure 1, right), where replicated samples were present, LDM-mode and LDM-median yielded higher sensitivity than LinDA, with an increase of up to 9%, which is attributable to our permutation-based inference. In general, whenever LinDA controlled the FDR, LDM-mode and LDM-median also controlled the FDR. When LinDA had inflated FDR, such as in S4C0 (Figures 1, left), S3C0 (Figure S7), and S6C0 (Figure S9), LDM-mode and LDM-median controlled the FDR or had less inflated FDR. The only exception is in S3C0 (Figure S7), where LDM-median had a more inflated FDR than LinDA. With sparse signals, use of the median for 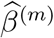 exhibited better sensitivity than use of the mode. Conversely, with dense signals, the mode-based estimator outperformed the median-based estimator. However, in both cases, using the median led to better power for community-level tests than using the mode (Figure S12).

**Figure 1:**
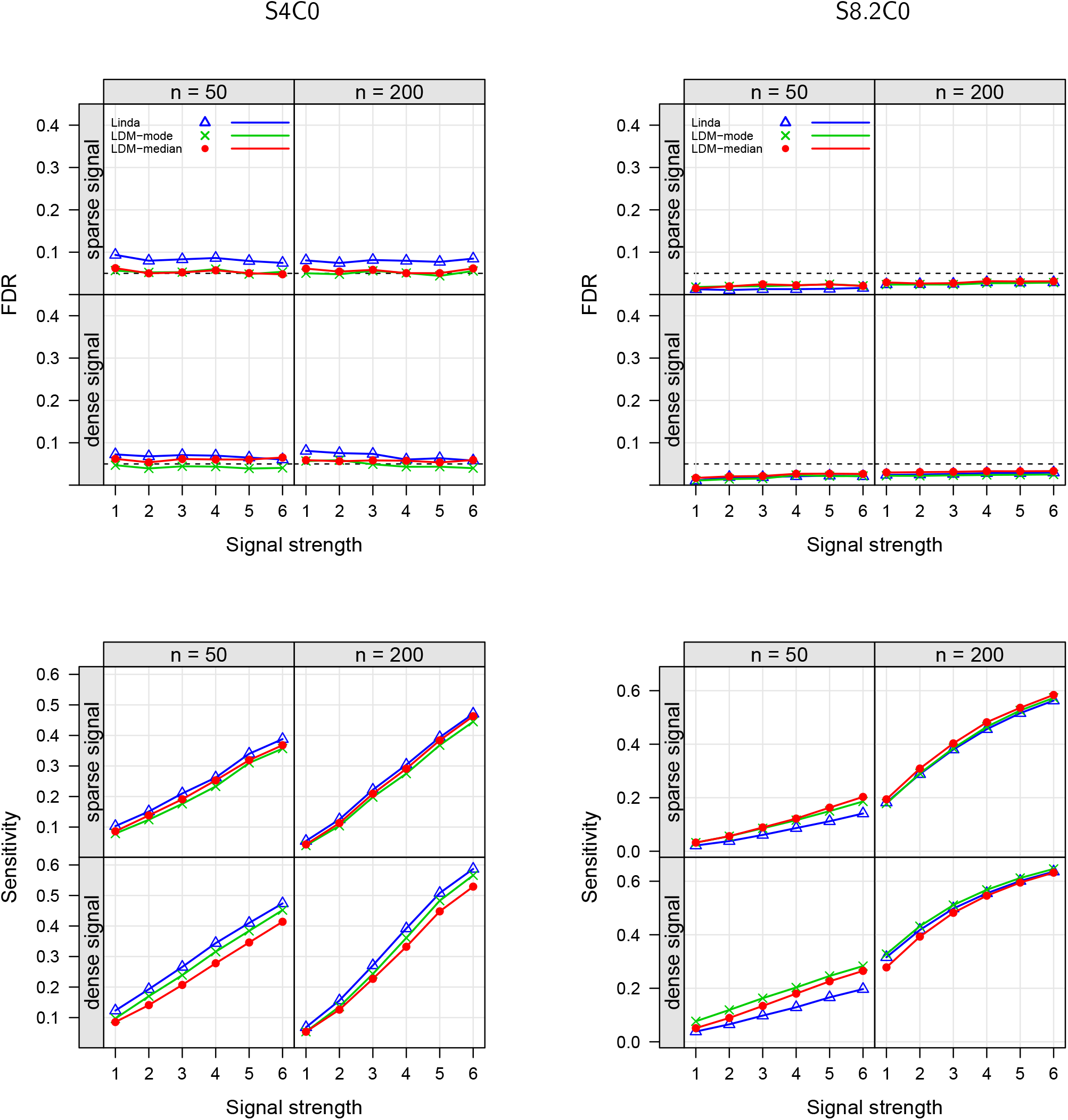
Empirical FDR and sensitivity at the nominal FDR level of 0.05 (black dashed line) in the S4C0 setting (with a small number of taxa) and the S8.2C0 setting (with replicated sampling). All results here as well as in other figures were based on 1000 replicates of data.

### Compositional analysis of the aGVHD data

We analyzed the 16S rRNA sequencing data on acute graft-versus-host disease (aGVHD) [18], which originally consisted 2436 operational taxonomic unites (OTUs) across 94 subjects. Following the guidelines in [9], we removed subjects with library sizes less than 1000 and excluded OTUs found in fewer than 10% subjects, resulting in a final dataset comprising 89 subjects and 304 OTUs for our analysis. We tested the association of the gut microbiome with two survival outcomes separately: the overall survival and the time to stage-III aGVHD, with censoring rates of 52.3% and 42.0%, respectively. Both analyses were adjusted for age and gender.

We previously established in [7] that censored survival times can be tested in association with microbiome data using LDM by treating the Martingale or deviance residual obtained from the Cox model as a continuous trait; the Cox model incorporates potentially confounding variables (such as age and gender in this case) as covariates, while excluding the microbiome data. LDM also provides a combination test that integrates the results from analyzing the two residuals. LDM-clr inherits these features, enabling testing of clr-transformed data with censored survival times at both taxon and community levels. Using the same approach, we can also apply LinDA to test censored survival times by manually calculating the Martingale or Deviance residual for these data, and then treating it as a covariate. However, LinDA does not support combination tests or community-level tests.

The results of our analyses are summarized in Table 1. For the time to stage-III aGVHD, LDM-mode detected 13 associated OTUs at the nominal FDR level of 5%, while LinDA detected 16, using the Martingale residual. Both methods detected 0 associated OTU using the Deviance residual, demonstrating a significant variability in results when choosing which one to use *a priori*. Fortunately, LDM-mode offers a combination test that detected 8 associated OTUs. The community-level *p*-values generated by both LDM-median and LDM-mode were all small, aligning with the detection of associated OTUs. For the overall survival, LDM-mode detected 3 OTUs, consistent with its small community-level *p*-value, while LinDA detected none.

**Table 1.**
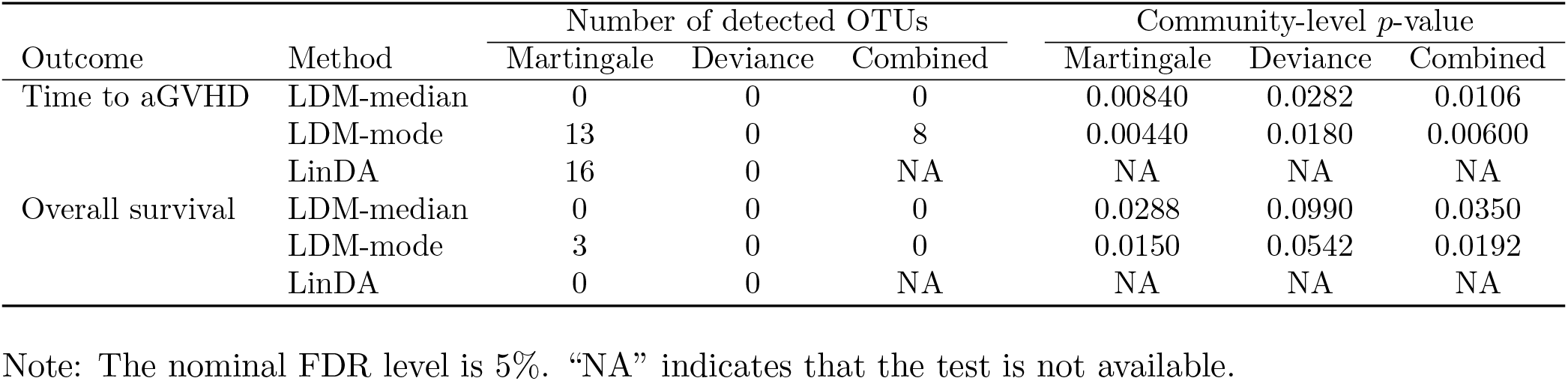
Results from compositional analysis of the aGVHD data

| Outcome | Method | Number of detected OTUs | | | Community-level $p$-value | | |
| --- | --- | --- | --- | --- | --- | --- | --- |
|  |  | Martingale | Deviance | Combined | Martingale | Deviance | Combined |
| Time to aGVHD | LDM-median | 0 | 0 | 0 | 0.00840 | 0.0282 | 0.0106 |
|  | LDM-mode | 13 | 0 | 8 | 0.00440 | 0.0180 | 0.00600 |
|  | LinDA | 16 | 0 | NA | NA | NA | NA |
| Overall survival | LDM-median | 0 | 0 | 0 | 0.0288 | 0.0990 | 0.0350 |
|  | LDM-mode | 3 | 0 | 0 | 0.0150 | 0.0542 | 0.0192 |
|  | LinDA | 0 | 0 | NA | NA | NA | NA |

Note: The nominal FDR level is 5%. “NA” indicates that the test is not available.

## Discussion

Recently, Zhou et al. [9] have proposed LinDA, a linear model for clr-transformed data that has some similarities with LDM-clr. One major difference between LinDA and the LDM family is that LinDA uses asymptotic distributional assumptions, so that *p*-values are calculated using normal (or *t* distributions in the case of small sample sizes) for independent samples; inference in LDM is permutation-based. Additionally, for correlated samples, LinDA uses asymptotic results for the linear mixed-effect model, which may fail if the distributional assumptions do not hold; by contrast, LDM-clr handles sample clustering by a distribution-free approach [4]. Further, inference in LinDA does not account for variability in 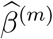 which we find is important when the number of taxa is small.

Another difference between LinDA and LDM is that LDM provides a unified framework of taxon-level and community-level tests, ensuring coherence between their results; LinDA does not support community-level tests. Specifically, the test statistic for the community-level test is a ratio, with numerator and denominator given by the sum of the taxon-specific *F* -statistic numerators and denominators, respectively. When the number of taxa *J* is large enough that the variability in 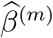 can be ignored, then the LDM-clr global test statistic is equal to the PERMANOVA test statistic based on Aitchison’s distance (the Euclidean distance of clr-transformed data) applied to data 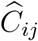 (defined just after (3)). Here we note that the Freedman-Lane permutation scheme used by the LDM and PERMANOVA-FL functions in the R package LDM actually gives slightly higher power than that attained by the implementation of PERMANOVA in the R package vegan. Even when the variability in 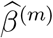 is accounted for, LDM-clr still produces test statistics that are similar to PERMANOVA based on 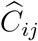 We note that comparison of LDM-clr and PERMANOVA using Aitchison’s distance is appropriate as this distance is considered the appropriate metric for community-level tests in compositional analysis of microbiome data [19]. Thus, we are confident that LDM-clr provides a useful addition to the growing list of methods for compositional analysis of complex microbiome data.

## Funding

This research was supported by the National Institutes of Health award R01GM141074 (Hu, Satten).

## Notes

### Competing Interest Statement

The authors have declared no competing interest.

