## Supplementary Materials for "Compositional analysis of microbiome data using the linear decomposition model (LDM)"

### Supplementary Figures

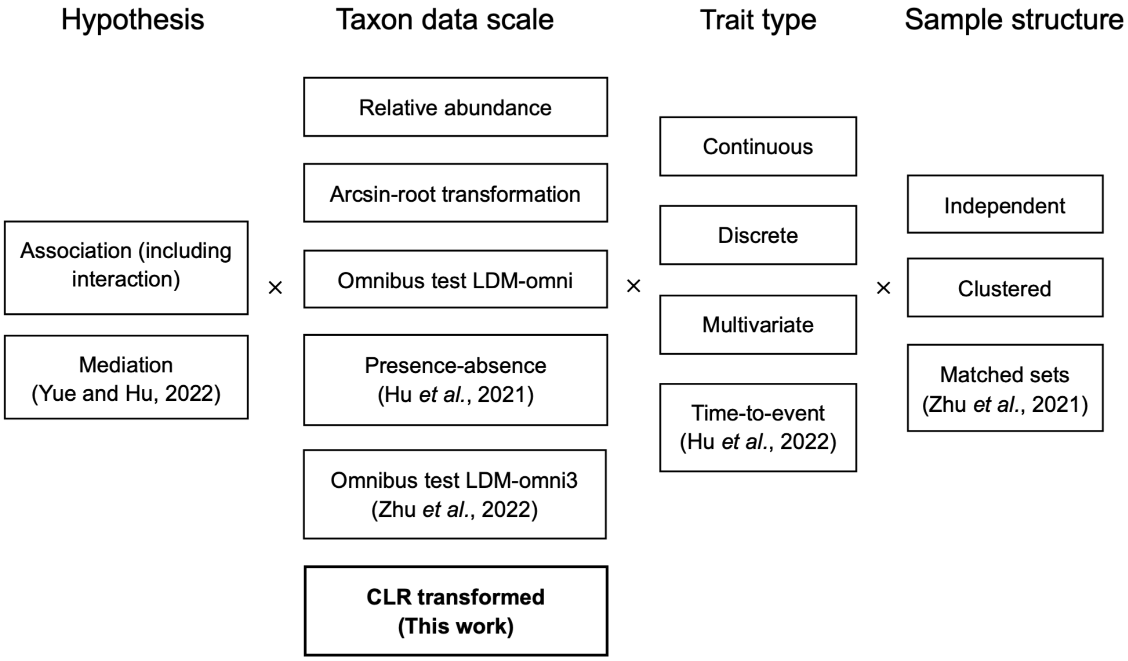


Figure S1: Analyses now supported by LDM. Analysis types without a citation were introduced in the original LDM paper [3]. "Clustered" refers to analyses of clustered data where traits of interest vary by cluster or vary both by and within clusters (referred to as "replicate sampling" in [9]). "Matched sets" is a special type of clustered analysis in which all traits of interest vary within sets (referred to as "pre-treatment" and "post-treatment" comparison in [9]).

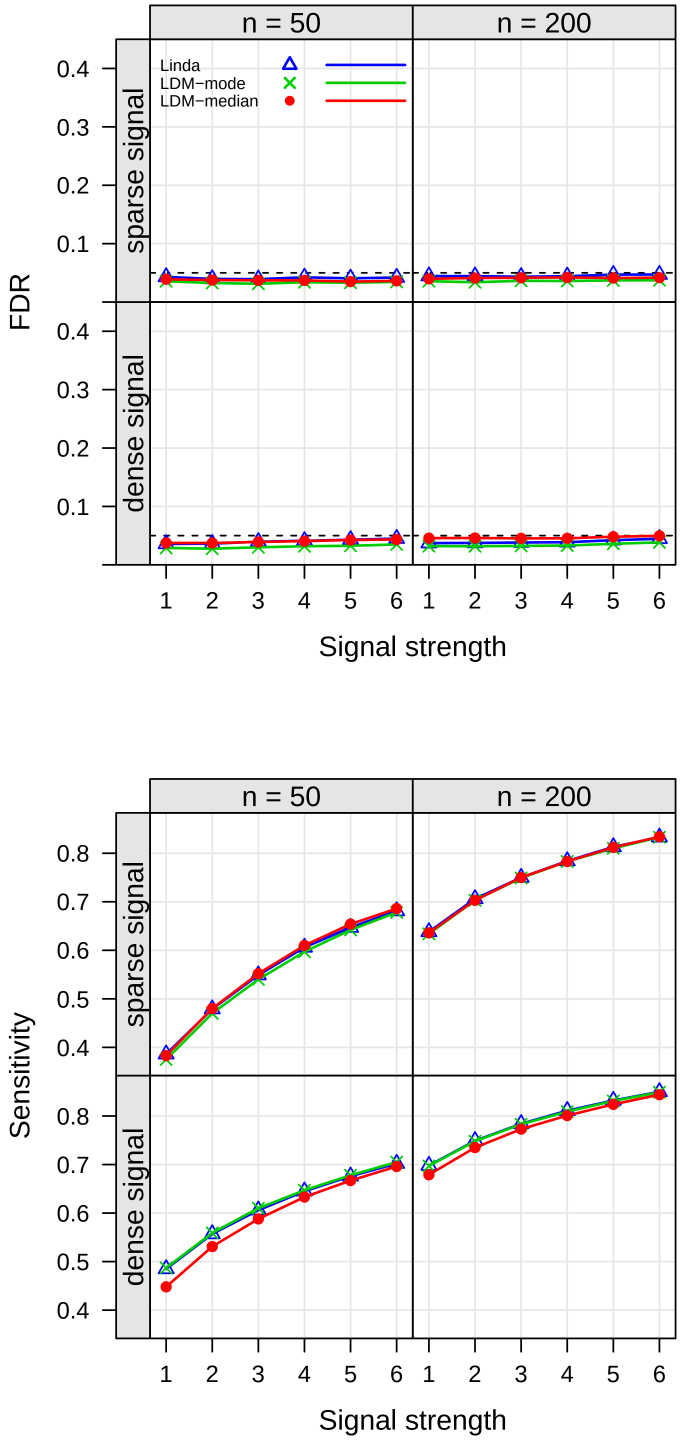


Figure S2: Simulation results for the setting with a binary trait (S0C0).

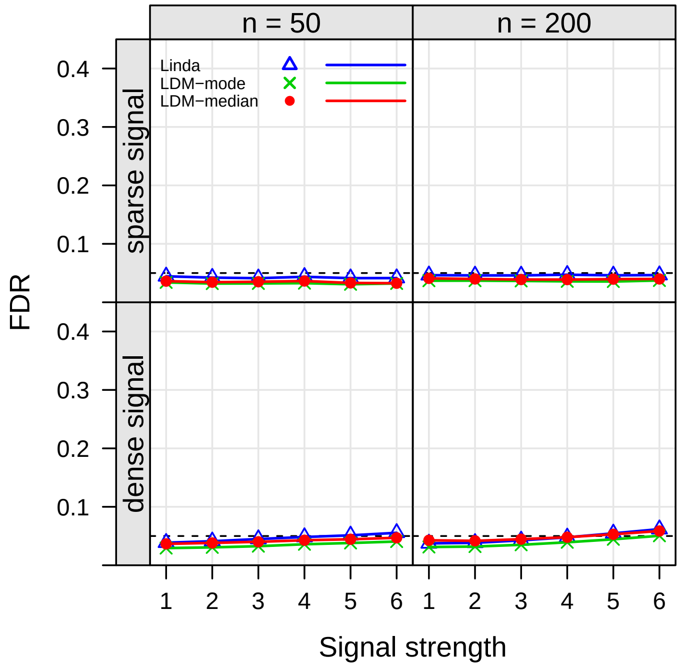


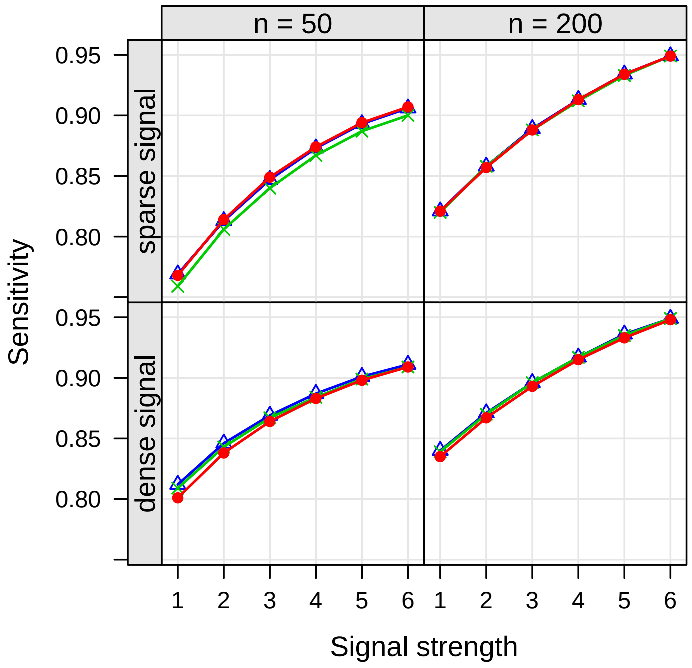


Figure S3: Simulation results for the setting with a continuous trait (S0C1).

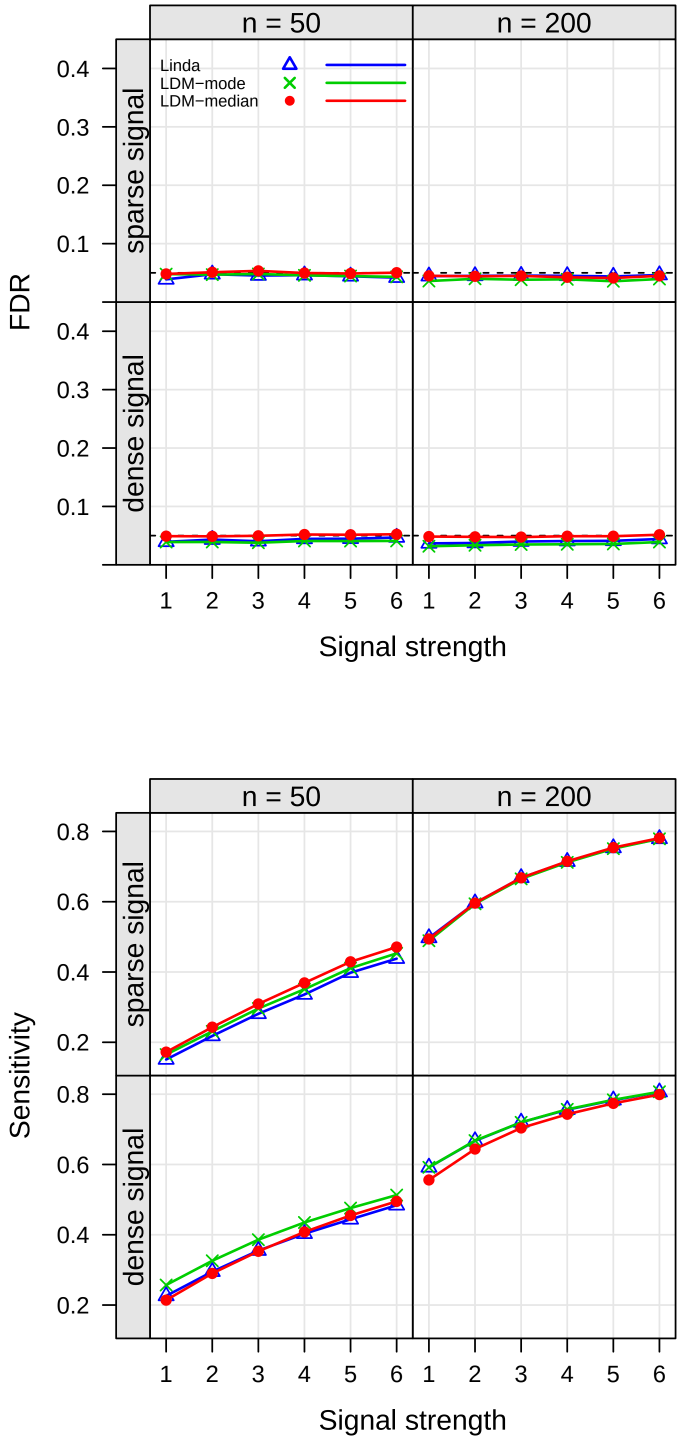


Figure S4: Simulation results for the setting with a binary trait and two confounders (S0C2).

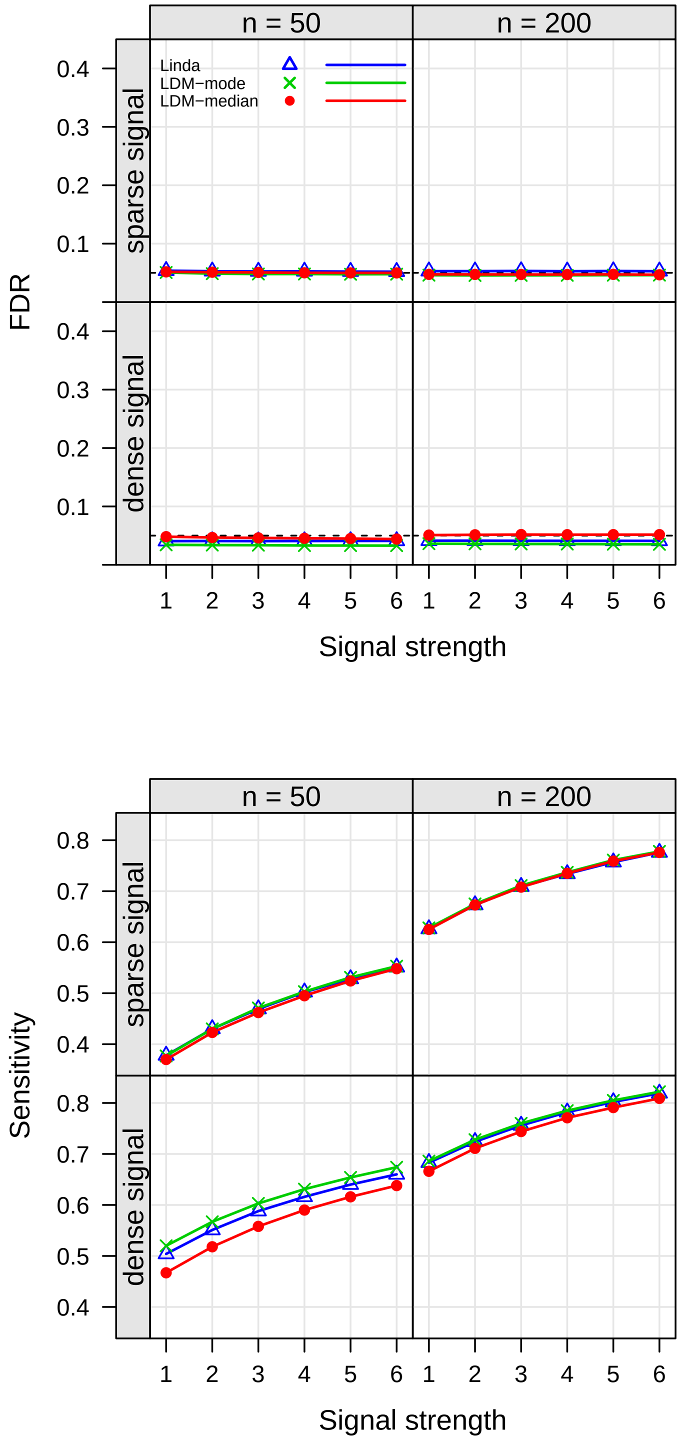


Figure S5: Simulation results for the setting with zero-inflated absolute abundances (S1C0).

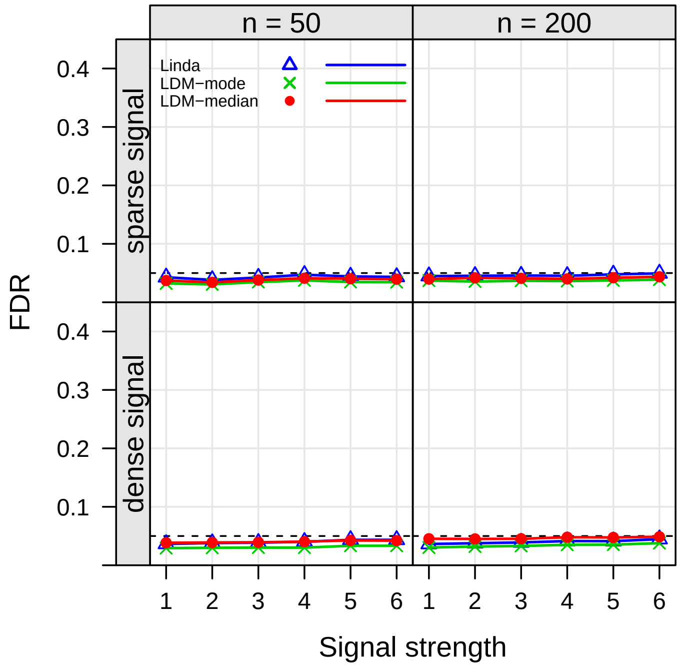


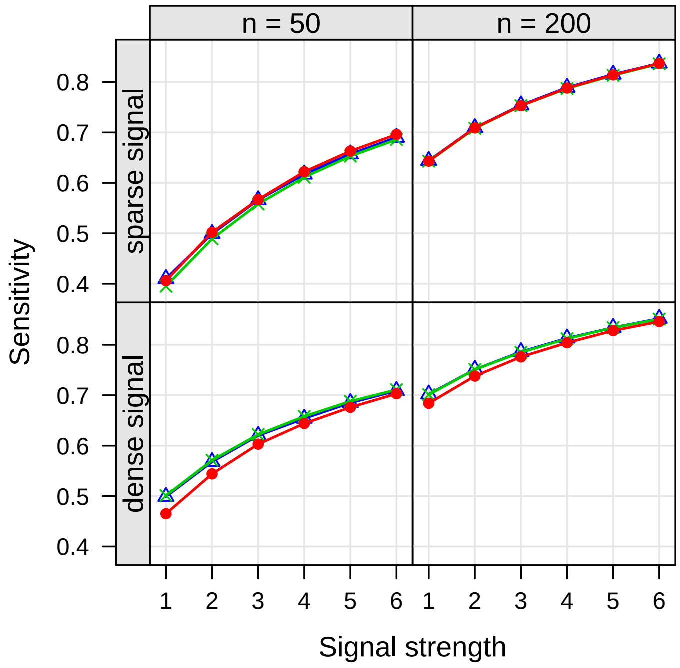


Figure S6: Simulation results for the setting with correlated absolute abundances (S2C0).

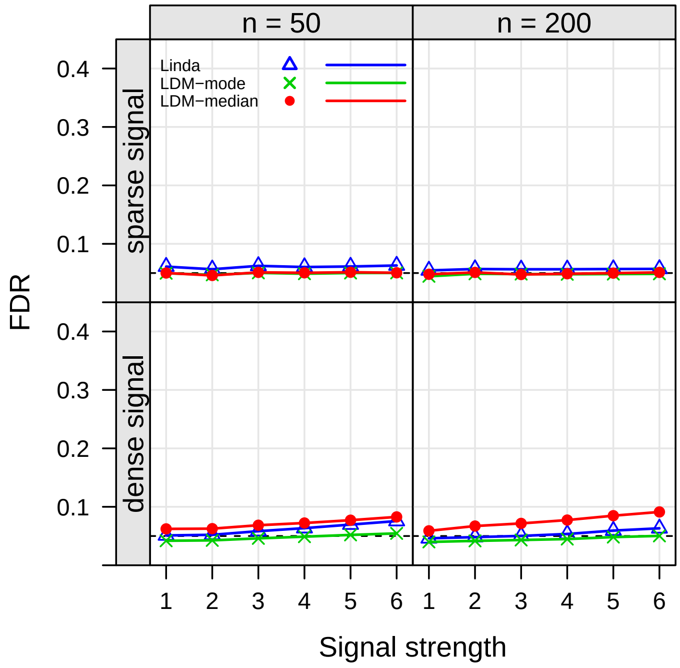


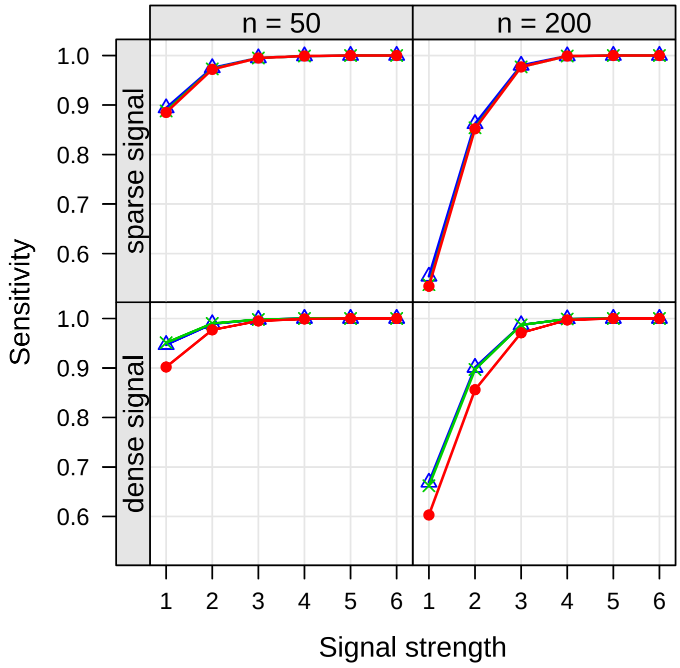


Figure S7: Simulation results for the setting with Gamma abundance distribution (S3C0).

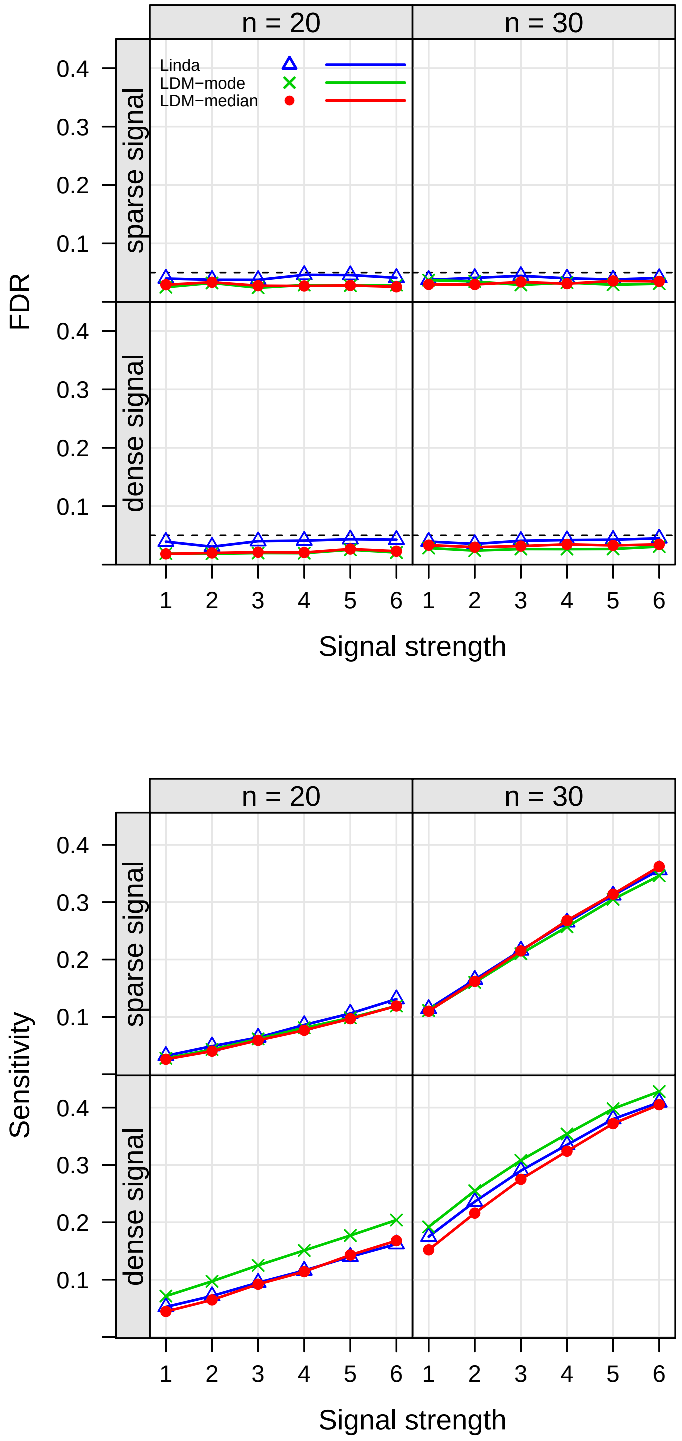


Figure S8: Simulation results for the setting with smaller sample sizes (S5C0).

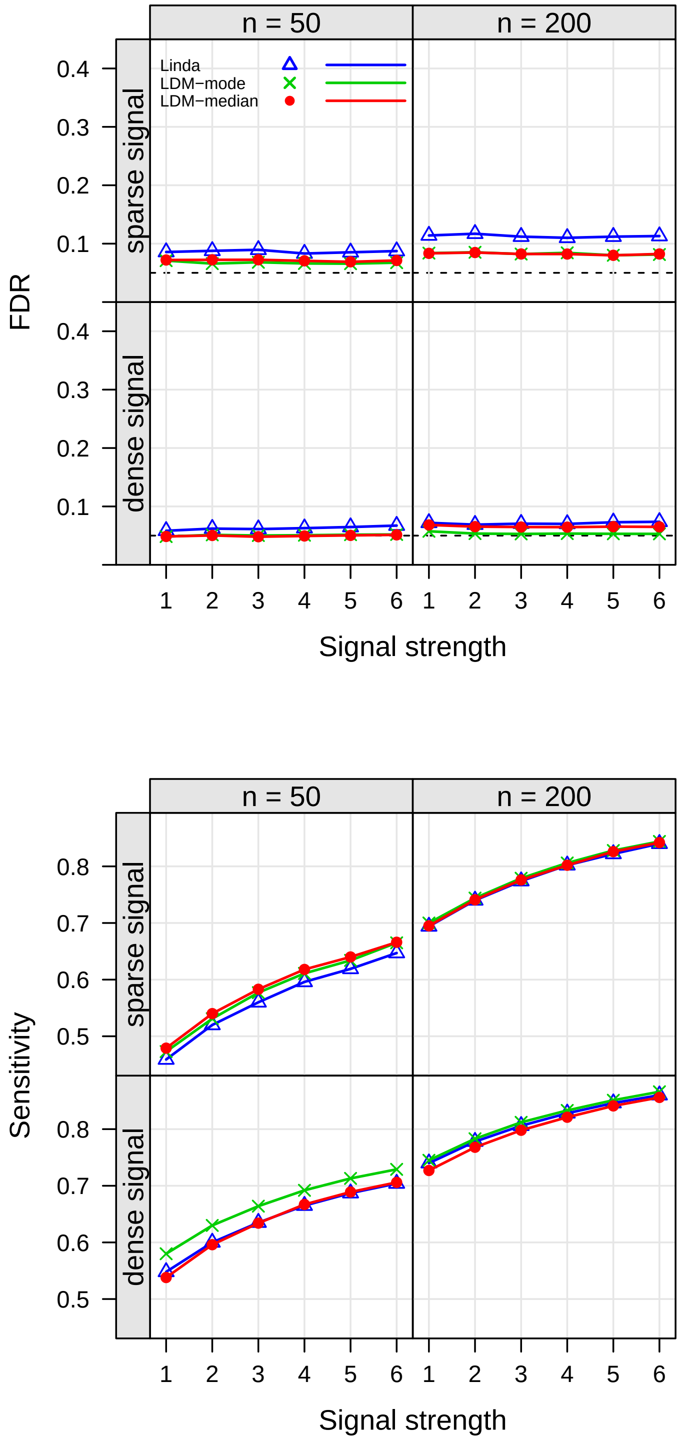


Figure S9: Simulation results for the setting with 10-fold difference in library size (S6C0).

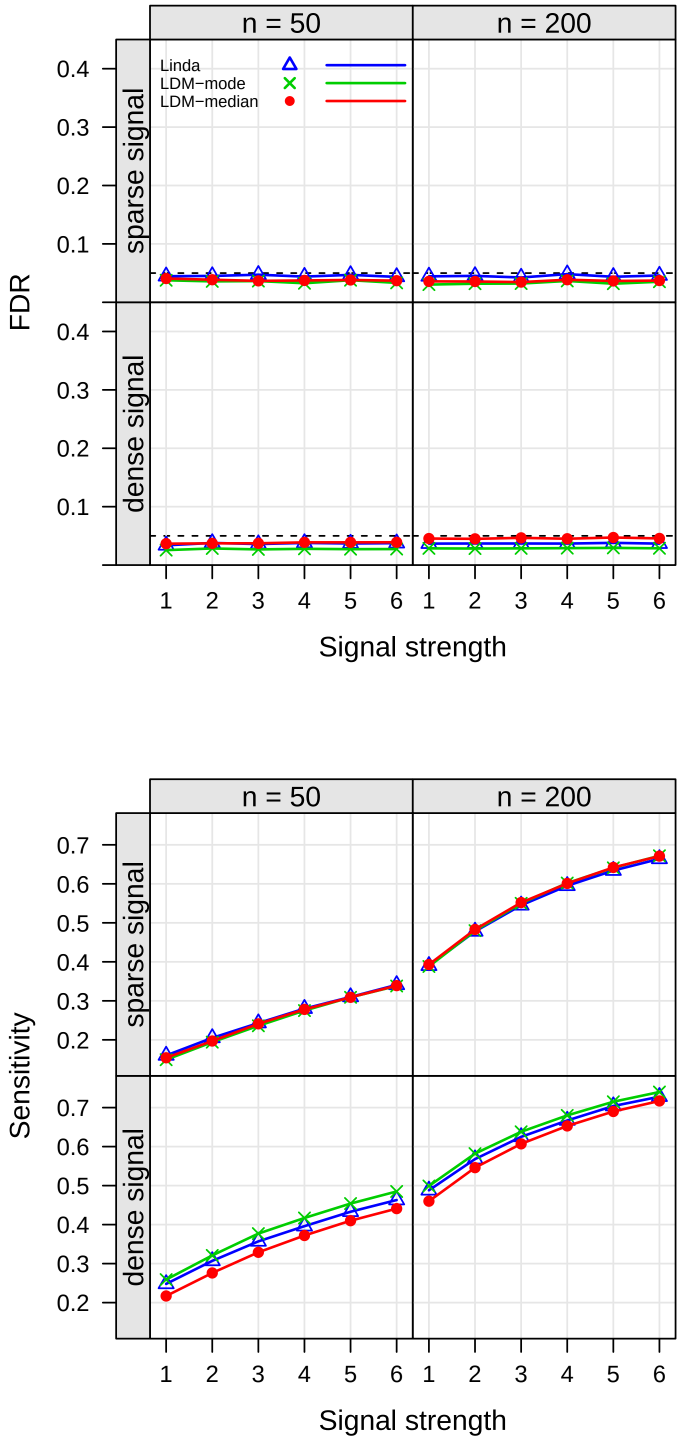


Figure S10: Simulation results for the setting with Negative-Binomial abundance distribution (S7C0).

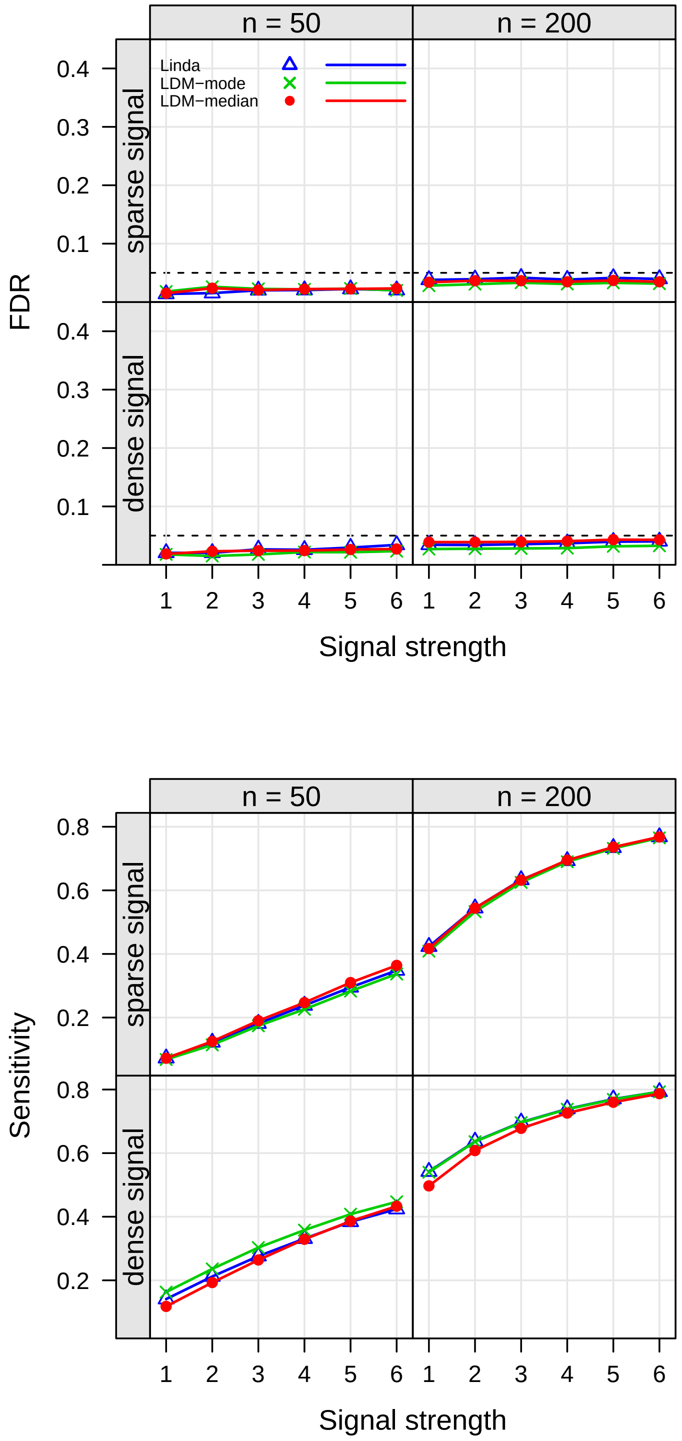


Figure S11: Simulation results for the setting with matched-set data (S8.1C0).

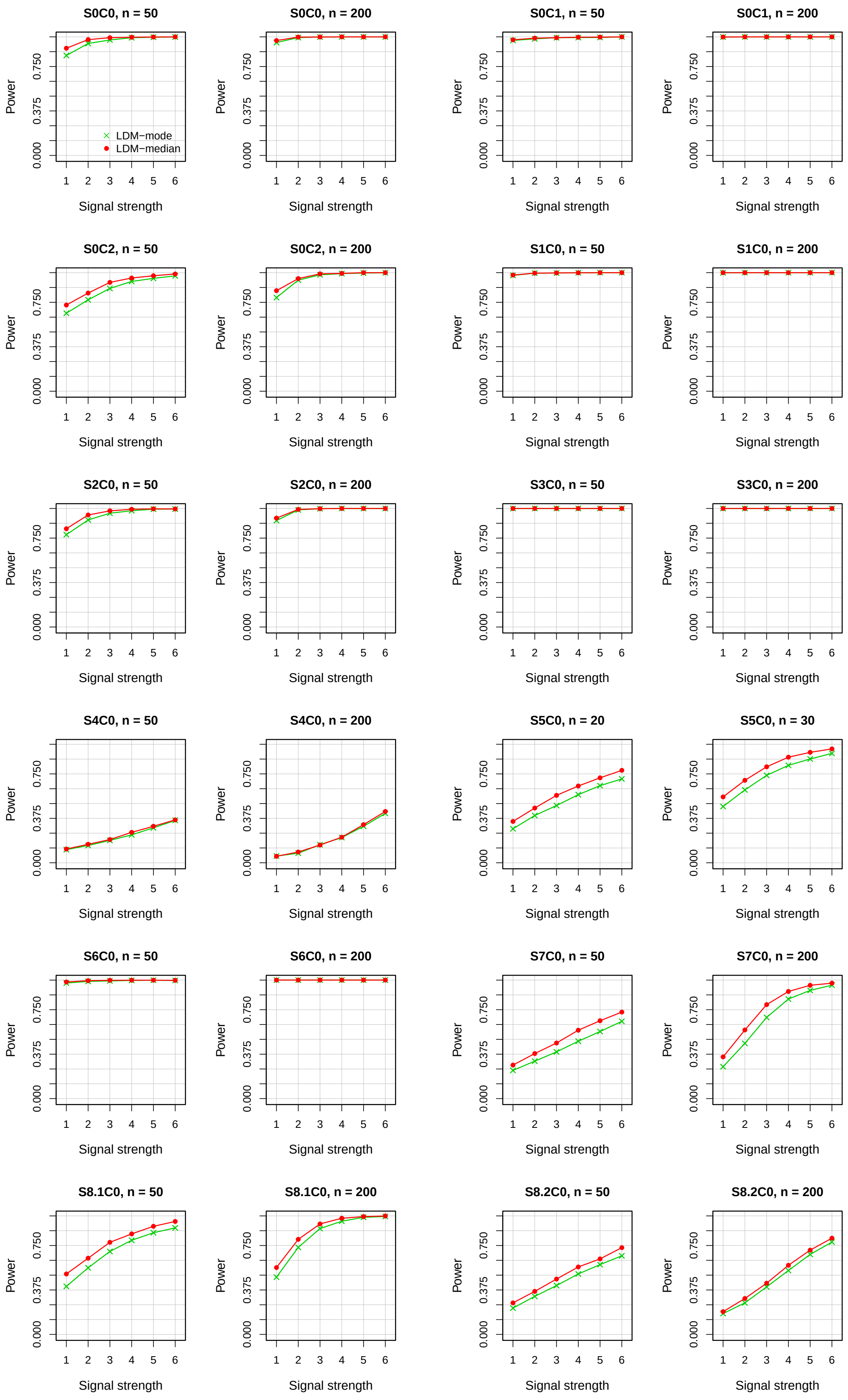


Figure S12: Power of the community-level tests at the nominal level of 0.05. All results were based on the "sparse signal" setting and 1000 replicates of data. Results from the "dense signal" setting are not shown because the power results are mostly 1 or, in a few cases when they are lower than 1, LDM-median is always slightly more powerful than LDM-mode.
